# Human-Brain Artificial-Intelligence Matrix

**DOI:** 10.1101/2020.09.09.288399

**Authors:** John Ibrahim

## Abstract

Human-Brain Artificial-Intelligence Matrix is a new technology aims to connect the human brain with the machine for the purpose of enabling the human brain to perform defined functions even if it becomes unable to perform them such as performing the function of vision in case of blindness, the function of hearing in case of deafness, Performing the function of motion in case of paralysis and many other functions. This technology will be based on the Cognition Theory which I argue about that the whole process of cognition can be treated quantum-mechanically. The cognition starts when a neuron sends data to be processed in the brain and ends in an effector to respond. The data “action potential” is a current of particles which can be described quantum-mechanically as a wave-impulse based on the dual nature of the particles. The neurons are a net of entangled cells classically and quantum-mechanically. When the action potential changes the potential of the neurons, it creates quantum mechanical potential wells and barriers. The action potential perfectly transmits in and out the neurons through quantum mechanical tunnels. The form of energy before processing is not the same after, but the amount of energy is always conserved. Since the neurons are entangled during the action potential transmission, the brain and effector will be entangled during the action potential processing. The effector’s cognition of data must be a discrete cognition of single-valued data from its self-adjoint matrix which entangled with brain matrix.

## I. THE COGNITION THEORY

### A. The fundamental postulate

The action potential arises as two entangled impulses^1^, one in a sensory neuron, and the other in the entangled motor neuron^2^. Next, the state of entanglement between neurons will be replaced with a state of entanglement between the brain and the effector.

#### 1. The scientific justification

**FIG. 1.**
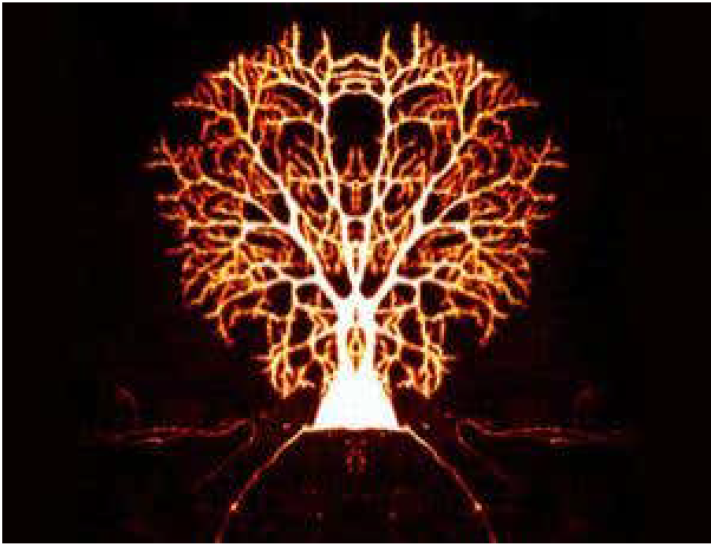
Entangled Neurons

Previous experiments have shown that the electrical activity of neurons in separate parts of the brain can oscillate simultaneously at the same frequency, a process known as phase locking. The frequency seems to be a signature that marks out neurons working on the same task, allowing them to identify each other. Now the observation that groups of brain cells seem to have their own version of quantum entanglement, could help explain how our minds combine experiences from many different senses into one memory”. Importantly, cloned signals only appeared after one region had reached a threshold level of activity. This threshold might ensure that our attention is only captured by significant stimuli rather than by every single signal^3,4^.

### B. The mathematical proof

#### 1. Pre-Cognition

Pre-Cognition is the process before the action potential transmitting to the brain.

1. The peripheral nervous system consists of neurons, Sensory and Motor which are quantum-mechanically entangled.
2. The action potential arises as two entangled impulses, one in a sensory neuron, and the other in the entangled motor neuron.
3. The action potential will be treated as a wave-impulse^5,6^: (*λ*), is the De Broglie wavelength of a particle where it can be found through dividing the Planck constant (*h*) by the particle’s momentum (*p*), or through multiplying the wave vector (*k*) by the particle’s momentum (*p*). Furthermore, The particle’s energy can be found through multiplying the Planck constant (*h*) by the particle’s angular frequency, (*ω*).

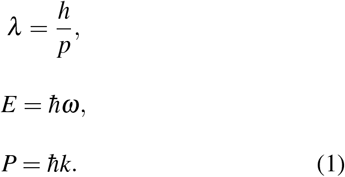 Treating the action potential as a wave-impulse will create a deep insight into how it transmits in and between neurons, and how the brain processes it.
4. The neurons will be treated as compound states of potential wells and barrier by the action potential mechanism.

By the thumb rule, a result could be found if the electric potential energy is growing or not. The electric properties of the action potential can be described by three main regions.

The first region is the depolarization state, where the potential difference increases from (−7m*mV*) to (0*mV*), which means the net potential difference is negative inside the membrane of the neuron, where the net charge is negative due to concentration of negative ions inside more than outside. The charge of the action potential is considered a flow of negative electrons. Hence, the electrical potential energy during this period is^7^:

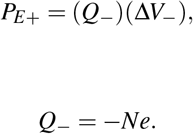

Hence, the potential energy is positive during this period which expressed quantum-mechanically as a potential barrier.

**FIG. 2.**
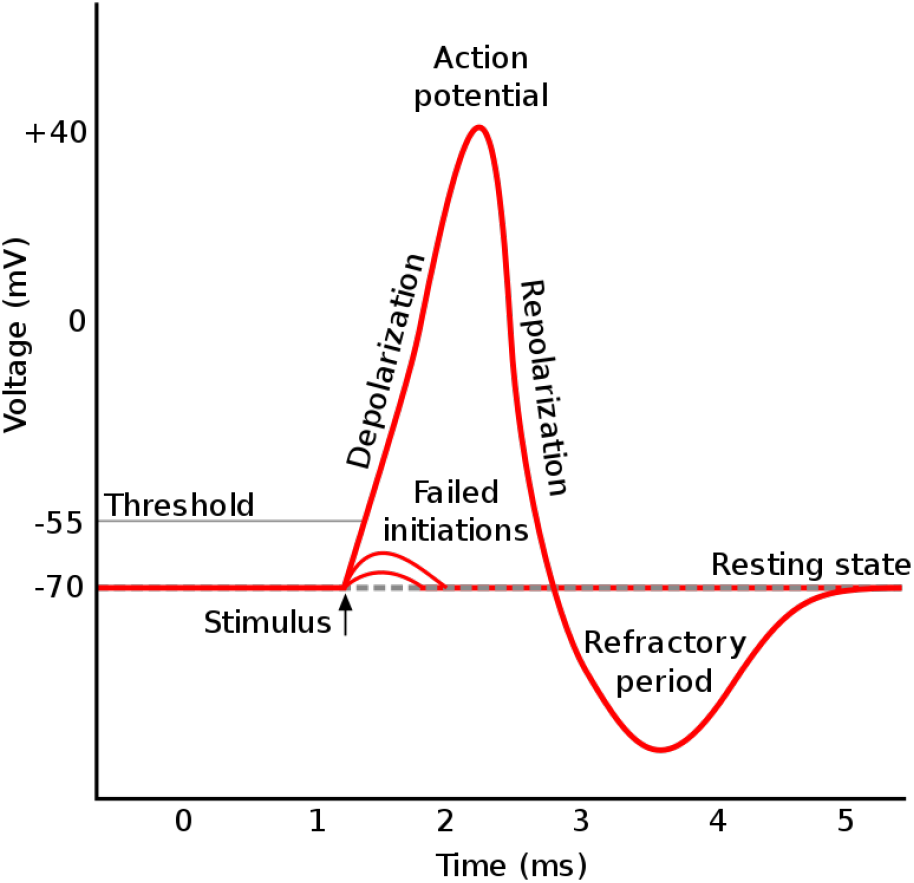
The action potential

The second region is a compound state of depolarization and re-polarization, from (0*mV*) to (40*mV*), then from (40*mV*) to (0*mV*), which means the net potential difference became positive inside the membrane of the neuron, where the net charge becomes positive due to flowing of the positive ions into the neuron’s membrane. The charge of the action potential is considered a flow of negative electrons. Hence, the electrical potential energy during this period is:

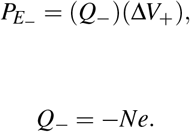

Hence, the potential energy is negative during this period which expressed quantum-mechanically as a potential well with a width [*a*].

The third region will be treated the same as the first, where the re-polarization is from(0*mV*) to (−70*mV*).

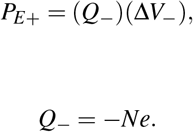

Hence, the potential energy is positive during this region which expressed quantum-mechanically a potential barrier.

That can explain clearly how the negative and positive ions oscillate inside and outside during the action potential. Where the exchange between the boundary quantum potential barriers and the middle well will allow the neuron to oscillate. Where the wave-function of the action potential can be written as:

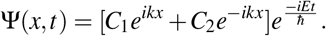

The crucial idea here is; how the impulse passes through the compound state of potential barrier-well-barrier, with a transmission coefficient equals one. That can be proved by considering a potential well with width (*a*) in the region (2), is between two potential barriers in regions (1) and (3). The energy of the impulse must be higher than the potential energy of the well and the barrier, (*E* > *V_b_* > *V_w_*), in order to avoid confining the action potential inside the potential well such as one-dimensional infinite well or finite well in quantum mechanics, when the energy of the electron is lower than the potential energy of the system^8^.

Generally, The Time Independent Schrödinger Equation:

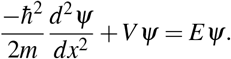

The Time Independent Schrödinger Equation [TISE] has different forms for each region:

Region 1: *x* < 0

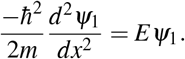

Region 2: 0 < *x* < *a*

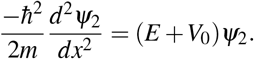

Region 3: *a* < *x*

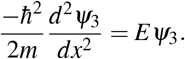

Solutions to the TISE in each region are, most generally:

Region 1: *x* < 0

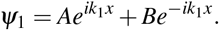

Region 2: 0 < *x* < *a*

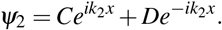

Region 3: *a* < *x*

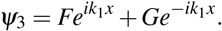

Where:

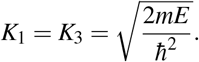

And,

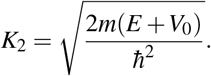

The derivatives of the functions are:

Region 1: *x* < 0

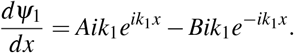

Region 2: 0 < *x* < *a*

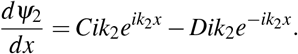

Region 3: *a* < *x*

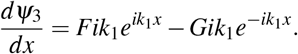

Since (*ψ*) and its derivative are finite, single-valued, and continuous, we can apply the following continuity relationships:

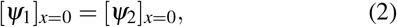

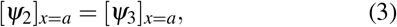

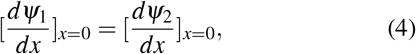

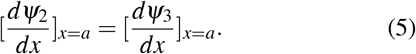

Since there should be no particles coming in from the far right moving in the negative x-direction, G must be zero.

Applying the continuity relationships on the equations (2), (3), (4), and (5), then, dividing all by (*i*), that gives:

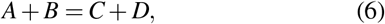

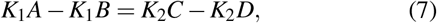

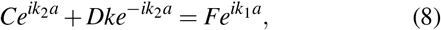

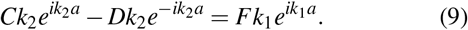

Thus equations (6), (7), (8), and (9) are the four necessary relationships between the five arbitrary constants A, B, C, D, and F. When rearranging equation (7):

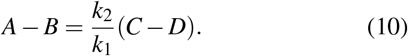

Then, adding equation (6) to (10):

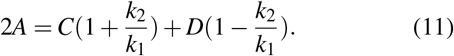

Then, rearranging equation (9):

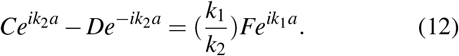

Then, adding equation (8) to equation (12), and subtract them after;

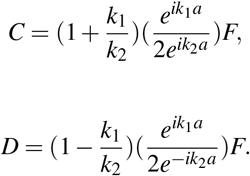

Then, substituting the value of C and D into equation (11):

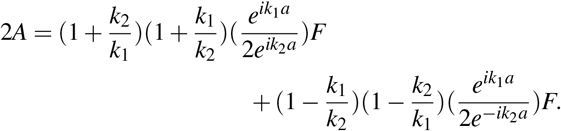

Then, factorising 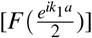:

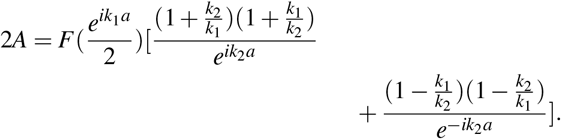

Then, rearranging to find the transmission coefficient 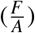:

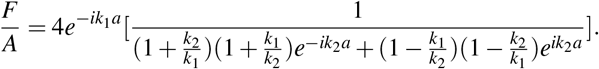

Then, expanding the brackets:

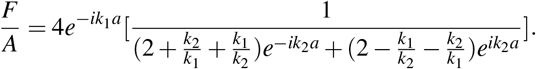

Then, taking (*k*_1_*k*_2_) as a common factor:

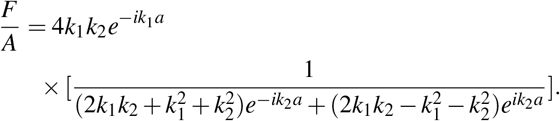

Then, expanding:

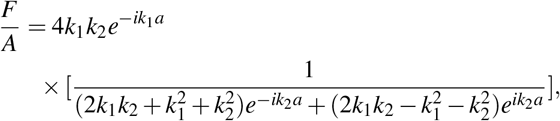

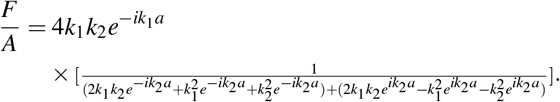

Then, rearranging:

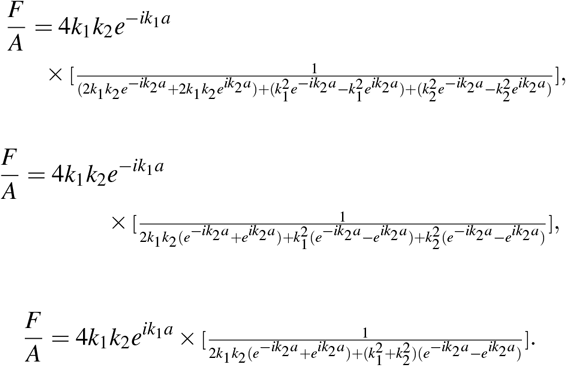

Using the complex trigonometric identities:

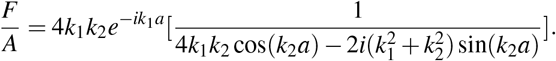

Simplifying the equation:

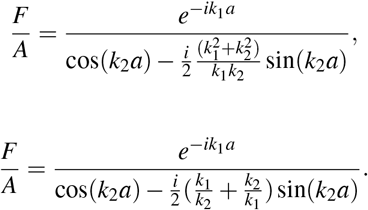

Then, multiplying by the complex conjugate:

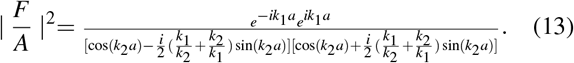

Finally, simplifying the equation (13):

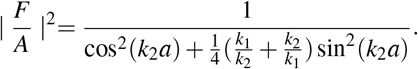

Hence, each particle of the action potential tunneling in the Sensory neurons will own this probability:

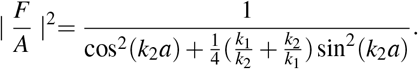

Now consider an entangled action potential, which will be arisen in the Motor neurons, tunneling in the opposite direction to the one in the Sensory neurons:

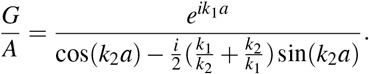

Then, multiplying by the complex conjugate:

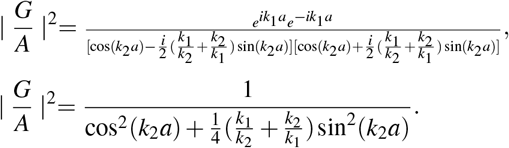

Hence, the particle which tunnel to the right will own this probability:

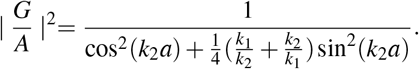

Therefore, both 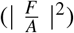 and 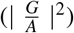 have the same exact probability for two particles; one will transmit from a side and the second will transmit from the opposite side.

Then, if considering the argument of cos^2^(*k*_2_*a*) and sin^2^(*k*_2_*a*), we find that two values (±*k*_2_*a*) are possible, since (cos) is an even function, and (sin) is squared.

Hence, I argue that one of these probabilities is related to the particle which emitted to the right-hand side and the other exact probability is related to the second particle which emitted to the left-hand side. Where the two particles when they are non-local entangled means they occupy two non-local states; one occupy a positive integer state and the other will occupy the same integer but with a negative sign. Where both will be produced with the same exact probability since the sin^2^(*k*_2_*a*) gives the same result for both positive and negative arguments since sine function is squared.

This deep insight is a prediction of the existence of the negative integer quantum states. Hence. the perfect transmission is done in a special case, where (*k*_2_*a* = *nπ*), if and only if (*n*) is an integer [n = −5, −4, −3, −2, −1,1,2,3,4,5].

Hence, the probabilities of tunneling are:

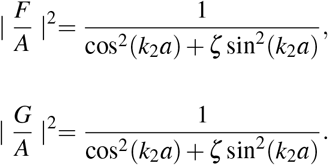

Where [*ζ*] zeta factor equals 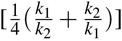

Now, finding the energy of the transmitted particle by considering any of the transmission coefficients; let it be 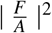:

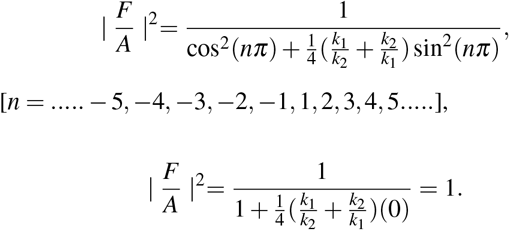

Considering the perfect transmission, by equating the two formulas of (*k*_2_):

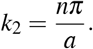

The case of (*n* = 0) is rejected depending on equation (13); (*k*_2_ = 0), which leads to divergence of the denominator.

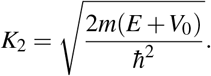

Hence, a formula for the allowed energies can be found;

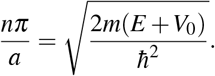

By squaring both sides:

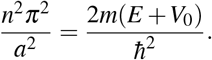

Finally, rearranging for (E) being the subject:

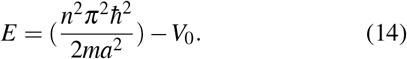

Equation (14) shows that the energies are quantized by (*n*^2^) factor which leads to the degeneracy of two states which means that two different states have the same energy; (*E*_(n=1)_ = *E*_(n=−1)_), (*E*_(n=2)_ = *E*_(n=−2)_), … and so on.

The deep quantum mechanics insight is the negative quantum states are a reflection of the positive quantum states on an axis which plays the role of the mirror that will induce arising a new higher dimension. Where both of them are non-local and have the same exact probability which means they are quantum-mechanically entangled. Therefore, I argue that the Sensory and the Motor neurons are entangled due to arising two entangled action potentials one in the Sensory neurons and one in the Motor neurons simultaneously.

Finally, equation (14) can explain clearly, “How can neurons distinguish between the different stimuluses ?”. Where the neurons will sort each specific stimulus to be qunatised uniquely in a specific quantised level.

#### 2. Post-Cognition

“Post-Cognition is the process after the action potential transmitting to the brain”

5-The total energy of the impulse before processing in the brain will be modelled as a Hamiltonian operator acting on the wave-function of the wave-impulse^9^.

When generalising equation (14):

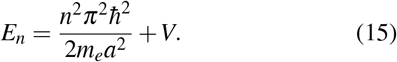

Where (*V*) is (−*V*_0_), the potential well energy at the brain when its neurons start to be stimulated, as shown in Fig 3.

**FIG. 3.**
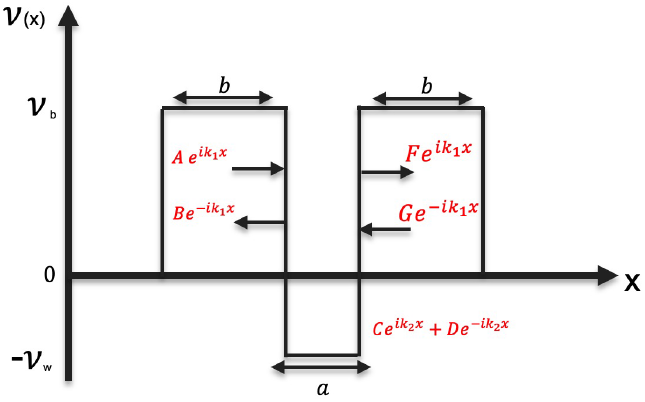
The compound state of potential barrier-well-barrier

Hence, the first term in the right-hand side represents the kinetic energy for each particle of the impulse, which can be written as

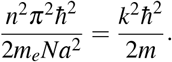

The second term in the left-hand side represents the potential energy for each particle of the impulse, which in one dimension can be written as:

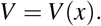

Then, rewriting equation (15) as:

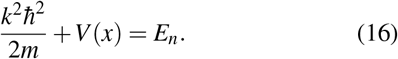

Using equation (1), the momentum of the impulse 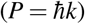 and squaring it;

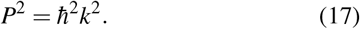

Substituting (17) in equation (16):

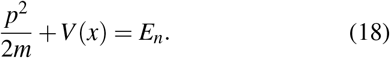

Then, finding the quantum mechanical operator of the momentum: 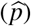

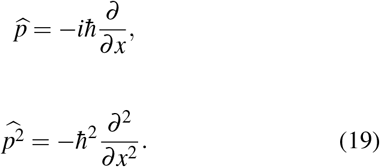

Substituting (19) in equation (18):

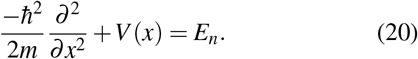

It is clearly shown that the left-hand side of equation (20) is the Hamiltonian operator 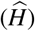

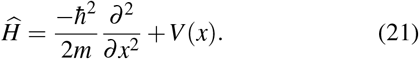

6-The brain activity will be treated as a Hermitian operator action on the impulse-wave creating processed data as discrete eigenfunctions. One of the most important postulates of quantum mechanics is: “Each physical observable is combined with a linear Hermitian operator; an eigenvalue of the operator is the measurement result. The result of [A]’s measurement is always one of the eigenvalues of [A]”

Depending on the previous results, since 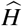, is a Hermitian operator. The equations (20) and (21) can be modified as:

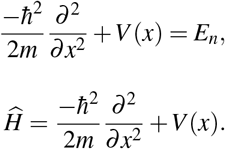

Considering that the wave-impulse has a wave-function, *ψ_n_*. Hence, the Hermitian operator (Hamiltonian) will act on the wave-function, *ψ_n_*, of the impulse as:

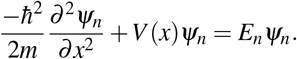

Hence, The equation can be modified in this way:

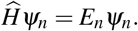

In quantum mechanics, when a Hermitian operator acts on the wavefunction, the outcomes always are the eigenvalues which are in this case the energies.

Depending on Sturm-Liouville theorem, the eigenvalues ‘energies’ can be considered a diagonal matrix of eigenvalues:

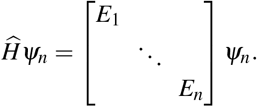

The result is that the brain processes arbitrary data through creation diagonal matrices of discrete eigenvalues.

7-The motor-entangled wave-impulse will be a self-adjoint carrying all the processed data to the effector, while the sensory impulse will be sent to the memory loop to be kept as a familiar pattern^10,11^. With applying [Sturm-Liouville Theory], that is a wide framework to some second-order ordinary differential equations. [Sturm-Liouville equation] in standard form is:

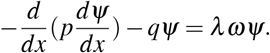

Same equation could be written as:

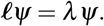

The Sturm-Liouville operator (ℓ) is:

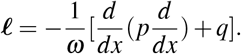

Applying a differential operator (*ℓ*) to a set of functions with (n= 1,2,...), then defining the following:

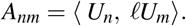

The values (*A_nm_*) with dissimilar [*n*] and [*m*] is represented as a matrix for the first index is the row while the second one is the column. The inner product definition says that if we reverse the arguments’ order, thus the result is changed to its complex conjugate.

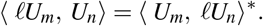

Let the operator (*ℓ*) and the boundary conditions with (*ℓ*) is self-adjoint, i.e., for all functions (*U*) and (*V*) with operator (*ℓ*):

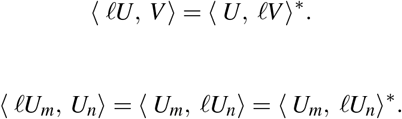

In the previous equation, in the first equality particularly, we have used the property of the self-adjoint. Then, in the second equality, exchanging the order of the arguments and taking the complex conjugate. Hence, the self-adjoint operator ensures that the matrix (*A*) will satisfy:

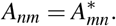

The complex conjugate of the matrix element is given by exchanging the two indices. Then, a matrix (*A*)’s transpose defined as (*A^T^*) can be obtained by interchanging the rows and columns of matrix (*A*), 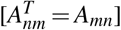. Hence, the complex conjugate of (*A*)’s transpose, 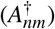, is the self-adjoint of a matrix (*A*),

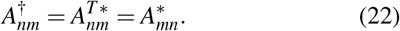

Using equation (22), it can be concluded that the matrix is said to be self-adjoint or Hermitian when it satisfies this condition, (*A*^†^ = *A*), which means that the matrix is equal to its own adjoint.

Considering the self-adjoint operator (*ℓ*)’s eigenfunctions (*U_n_*), due to the orthogonal property of these eigenfunctions, it can be concluded that,

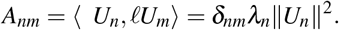

When these eigenfunctions are orthonormal, 〈*U_n_,U_m_*〉 = *δ_nm_*, that implies the matrix (*A*) is diagonal and that the matrix element (*Ann*) is equal to the relevant eigenvalue (*n*).

Hermitian operators in qunatum mechanics could be defiend as self-adjoints where the physical observables such as energy are corresponded to it. These physical observables are real where they present real eigenvalues. For these reasons, Schrödinger’s equation is regarded as special cases of the Sturm-Liouville equation. In our case, the physical observable is the energy which is a real eigenvalue. Hence we conclude two things:

1. The created processed matrix in the brain and the created self-adjoint matrix in the effector are identical.
2. The sensory entangled impulse will not be annihilated according to the conservation of energy law. Hence, it will be kept as a pattern in the brain, especially in the memory loop, which can explain how we remember the familiar matters and how we learn in pattern methodology^4^.

### 3. Results

The results obtained from the Cognition theory can be summarised as:

- The first, the peripheral nervous system consists of neurons; Sensory and Motor, which are quantum-mechanically entangled.
- The second, the action potential arises as two entangled impulses, one in a sensory neuron, and the other in the entangled motor neuron.
- The third, the action potential can be treated as a wave-impulse.
- The fourth, the neurons can be treated as compound states of potential wells and barriers by the action potential mechanism.
- The Fifth, the neurons sort out each specific stimulus to be qunatised uniquely in a specific quantised level.
- The sixth, the total energy of the impulse before processing in the brain can be modelled as a Hamiltonian operator acting on the wavefunction of the wave-impulse.
- The seventh, the brain activity will be treated as a Hermitian operator action on the impulse-wave creating processed data as discrete eigenfunctions.
- The eighth, the motor-entangled wave-impulse is a self-adjoint carrying all the processed data to the effector, while the sensory impulse will be sent to the memory loop to be kept as a familiar pattern.

## II. THE PRACTICAL APPLICTION

### A. Outlines

The first, the second, the third, the fourth and the fifth result tell that the Sensory neurons and the Motor neurons can be replaced by a machine which is able to pick up the same stimulus and turn it into two entangle currents, an action potential in the language of neurons. Next, they will be transmitted directly, one to the brain and the other to the effector. Just then, the state of entanglement between neurons will be replaced with a state of entanglement between the brain and the effector^12 13 14^.

The sixth, the seventh and the eighth result tell us how the brain processes data, where this role in case of brain deficiency, can be replaced by a machine processing data by the same mechanism^10,11^.

### B. General application

#### 1. The entangled Sensory-Motor neurons deficiency

##### Degree 1

A machine which is able to pick up the same stimulus, turn it into two entangle action potentials and transmit them directly, one to the brain and the other to the effector. This machine will replace the role of the Sensory-Motor neurons and play three main roles. The first role is to pick up the stimulus for example in case of deficiency in the optic nerve, photo-electric cells will be used to pick up light. The second role is to turn this picked stimulus into two entangled impulses, where the machine will do this role will be by Single-electron transistor (SET)^15^, which induces a Cooper pair that allow a pair of electrons to flow freely in superconductors. The third role is to transmit the entangled action potentials directly, one to the brain and the other to the effector, where the machine will do this will micro-electronic components which will be implanted at two positions, the surface of brain and surface of the effector. The case of the entangled Sensory-Motor neurons deficiency will be referred to as Degree 1.

#### 2. The entangled Brain-Effector system deficiency

##### Degree 2

This case is more difficult than the previous one, hence, new roles will be added to the machine by using artificial intelligence. The machine will play the role of guidance where an artificial intelligence robot which able to process the data in the same way the brain does, will be entangled with both the effector and the brain will guide the new type of cells to perform an additional role to their natural role. These cells will be in the Frontal lobe, where the data will be processed by artificial intelligence robot as problem-solving and the final solution [the matrix] will be entangled with the action potential in the cells of the Frontal lobe, just then, these cells will do the additional role properly. The case of the entangled Brain-Effector system deficiency will be referred to as Degree 2.

## III. THE PROTOTYPE

### A. A prototype for Blindness of Degree 1

A blind of degree 1 has a deficiency in the entangled Sensory-Motor neurons^16^

#### 1. The components

1. A Photo-electric cell to upload the data from the absorbed photon on the free escaped electrons.
2. Single-electron transistors (SET)^15^, which induces a Cooper pair that allow a pair of electrons to flow freely in super-conductors. It is clear that we can include the photo-electric cell which is made up of a superconductor with the Single-electron transistor (SET) to be one simple electronic component.
3. Neuralink’s electronic probes which transmit data to the brain and the effector accurately^17^.

**FIG. 4.**
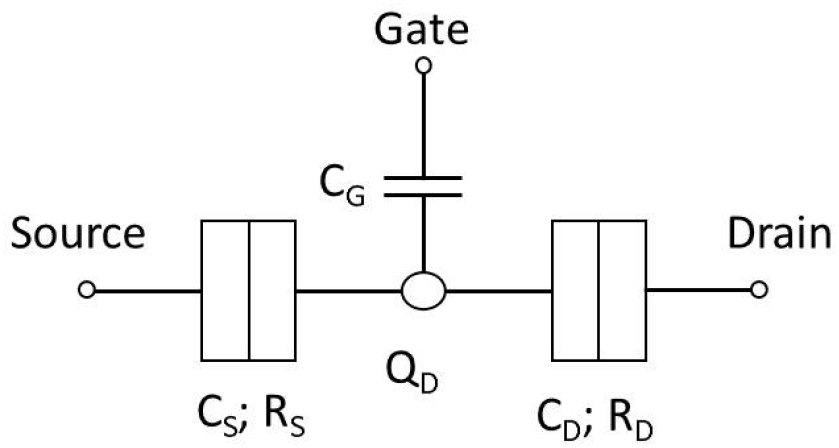
Single-electron transistor

**FIG. 5.**
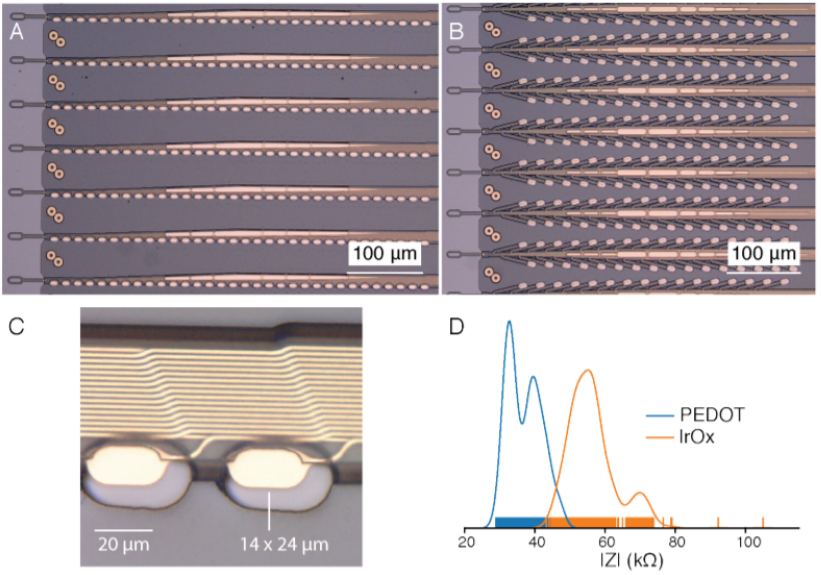
Neuralink’s electronic probes

#### 2. The mechanism

1. Two photoelectric cells are placed in front of the eyes as google glasses will be struck with photons of light.
2. These photons carry the visual data and will upload them on the electrons when striking them and allowing them to be free.
3. Four single-electron transistors (SET) which are connected; two at each end of the photo-electric cell, will induce a Cooper pair that allows a pair of electrons to flow freely in super-conductors.
4. On each side, two entangled currents will be created as two entangled action potentials, one will be transmitted to the brain by Neuralink’s electronic probe, which is implanted to be connected to the occipital lobe, and the second will be transmitted to the effector the Optic nerve in the same mechanism.
5. The brain will process the data as diagonal matrices of eigenvalues. The data in the optic nerve will be processed as self-adjoint matrices simultaneously. Just then, the blind of degree 1 will see.

**FIG. 6.**
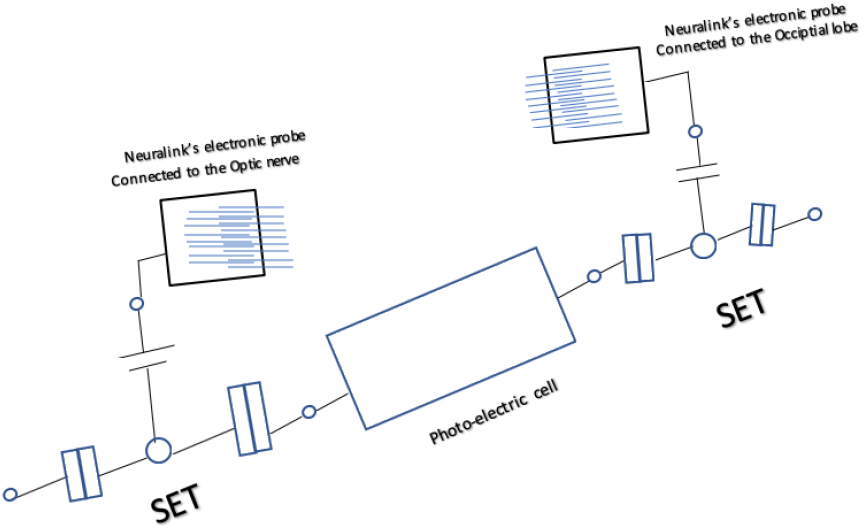
Prototype for Blindness of Degree 1

### B. A prototype for Blindness of Degree 2

#### 1. The components

1. A Photo-electric cell to upload the data from the absorbed photon on the free escaped electrons.
2. Single-electron transistors (SET) which induce a Cooper pair that allows a pair of electrons to flow freely in superconductors. It is clear that we can include the photo-electric cell which is made up of a superconductor with the Single-electron transistor (SET) to be one simple electronic component.
3. Neuralink’s electronic probes which transmit data to the brain and the effector accurately.
4. Artificial intelligence robot which is able to process the visual data as diagonal matrices of eigenvalues.

#### 2. The mechanism

1. Two photoelectric cells are placed in front of the eyes as google glasses will be struck with photons of light.
2. These photons carry the visual data and will upload them on the electrons when striking them and allowing them to be free.
3. Four single-electron transistors (SET) which are connected; two at each end of the photo-electric cells, will induce a Cooper pair that allows a pair of electrons to flow freely in superconductors.
4. On each side, two entangled currents will be created as two entangled action potentials, one will be transmitted to the brain by Neuralink’s electronic probe, which is implanted to be connected to the Frontal lobe, and the second will be transmitted to the Artificial intelligence robot, which can be connected directly to the Single-electron transistor (SET).
5. The Artificial intelligence robot will process the data as diagonal matrices of eigenvalues. Simultaneously, the data will be processed as final solutions; self-adjoint matrices, in Frontal lobe. Therefore, the cells of Frontal lobe will gain an additional role; the role of sight. Just then, the blind of degree 2 will see.

**FIG. 7.**
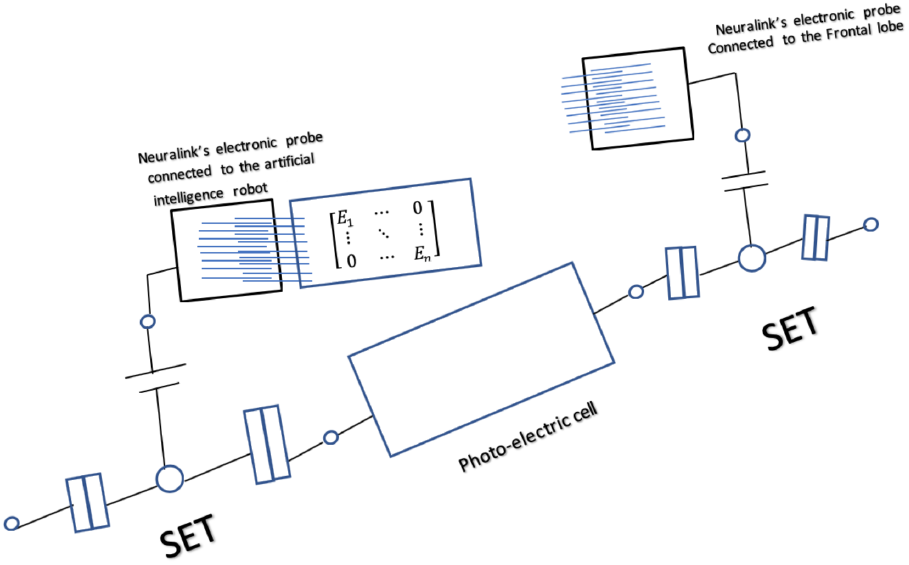
Prototype for Blindness of Degree 2

### C. A prototype for Deafness of Degree 1

A deaf of degree 2 has a deficiency in the entangled Brain-Effector system

#### 1. The components

1. Two small earphone cells.
2. Single-electron transistors (SET) which induce a Cooper pair that allows a pair of electrons to flow freely in superconductors. It is clear that we can include the small head-phone cells which are made up of a superconductor with the Single-electron transistor (SET) to be one simple electronic component.
3. Neuralink’s electronic probes which transmit data to the brain and the effector accurately.
4. Artificial intelligence robot which is able to process the auditory data as diagonal matrices of eigenvalues.

#### 2. The mechanism

1. Two small earphone cells to perform the function of generating electric-impulses as nerve impulses carrying auditory information.
2. The auditory data be uploaded on the generated electric impulses.
3. Four single-electron transistors (SET) which are connected; two at each end of the small earphone cells, will induce a Cooper pair that allows a pair of electrons to flow freely in superconductors.
4. On each side, two entangled currents will be created as two entangled action potentials, one will be transmitted to the brain by Neuralink’s electronic probe, which is implanted to be connected to the Frontal lobe, and the second will be transmitted to the Artificial intelligence robot, which can be connected directly to the Single-electron transistor (SET).
5. The Artificial intelligence robot will process the data as diagonal matrices of eigenvalues. Simultaneously, the data will be processed as final solutions; self-adjoint matrices, in Frontal lobe. Therefore, the cells of Frontal lobe will gain an additional role; the role of hearing. Just then, the deaf of degree 2 will hear.

**FIG. 8.**
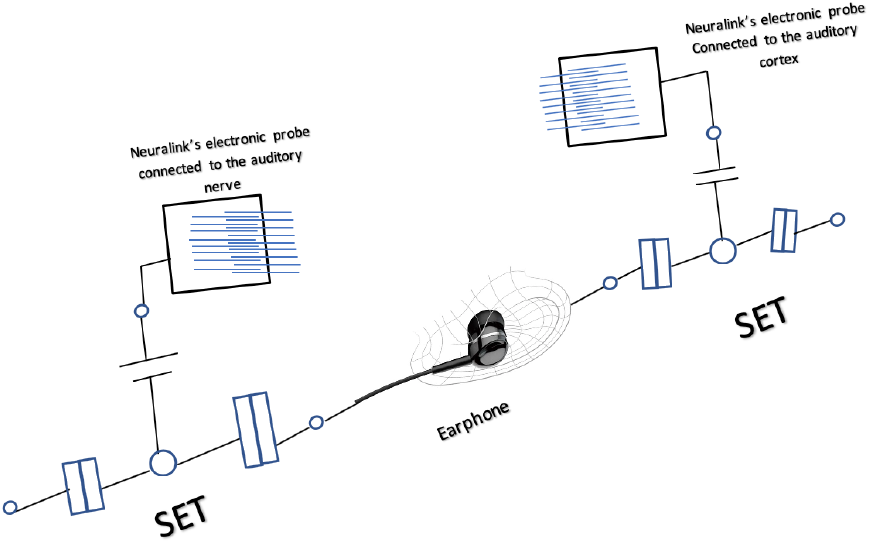
Prototype for Deafness of Degree 1

### D. A prototype for Deafness of Degree 2

A deaf of degree 2 has a deficiency in the entangled Brain-Effector system

#### 1. The components

1. Two small earphone cells.
2. Single-electron transistors (SET) which induce a Cooper pair that allows a pair of electrons to flow freely in superconductors. It is clear that we can include the small head-phone cells which are made up of a super-conductor with the Single-electron transistor (SET) to be one simple electronic component.
3. Neuralink’s electronic probes which transmit data to the brain and the effector accurately.
4. Artificial intelligence robot which is able to process the auditory data as diagonal matrices of eigenvalues.

#### 2. The mechanism

1. Two small earphone cells to perform the function of generating electric-impulses as nerve impulses carrying auditory information.
2. The auditory data be uploaded on the generated electric impulses.
3. Four single-electron transistors (SET) which are connected; two at each end of the small earphone cells, will induce a Cooper pair that allows a pair of electrons to flow freely in superconductors.
4. On each side, two entangled currents will be created as two entangled action potentials, one will be transmitted to the brain by Neuralink’s electronic probe, which is implanted to be connected to the Frontal lobe, and the second will be transmitted to the Artificial intelligence robot, which can be connected directly to the Single-electron transistor (SET).
5. The Artificial intelligence robot will process the data as diagonal matrices of eigenvalues. Simultaneously, the data will be processed as final solutions; self-adjoint matrices, in Frontal lobe. Therefore, the cells of Frontal lobe will gain an additional role; the role of hearing. Just then, the deaf of degree 2 will hear.

**FIG. 9.**
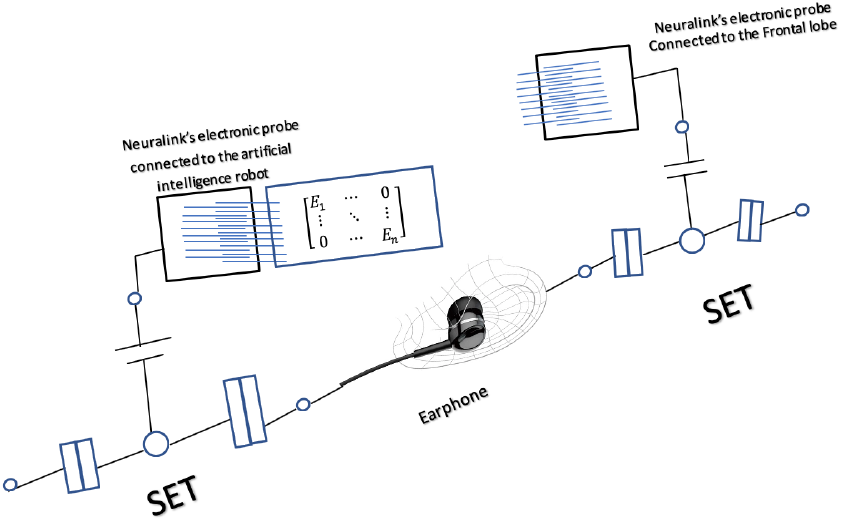
Prototype for Deafness of Degree 2

### E. A prototype for Paralysis of Degree 1

A paralysed of degree 1 has a deficiency in the entangled Sensory-Motor neurons^18^

#### 1. The components

1. Neuralink’s robotic electrode inserter.
2. Single-electron transistors (SET) which induce a Cooper pair that allows a pair of electrons to flow freely in superconductors. It is clear that we can include the single-electron transistor (SET) at the surface of the Neuralink’s robotic electrode inserter.
3. Neuralink’s electronic probes which transmit data to the brain and the effector accurately.

**FIG. 10.**
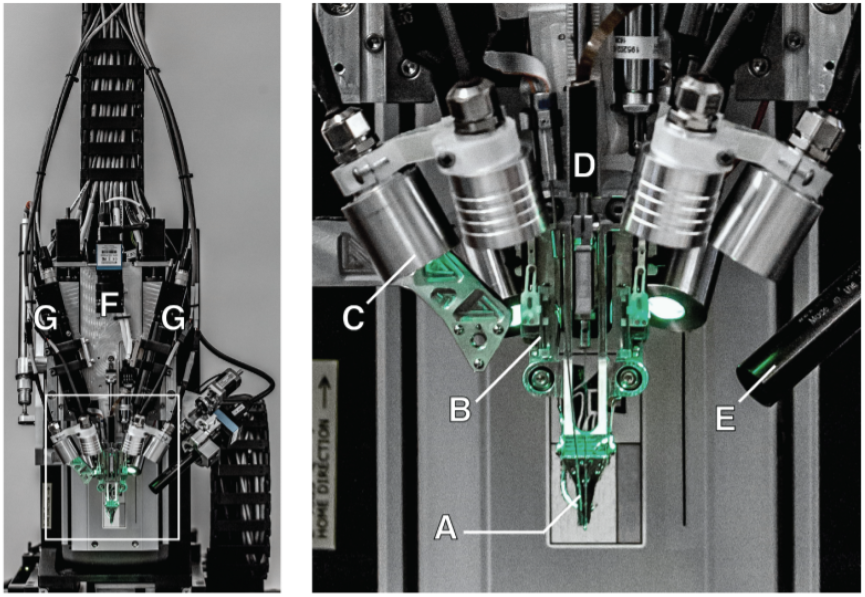
Neuralink’s robotic electrode inserter.

**FIG. 11.**
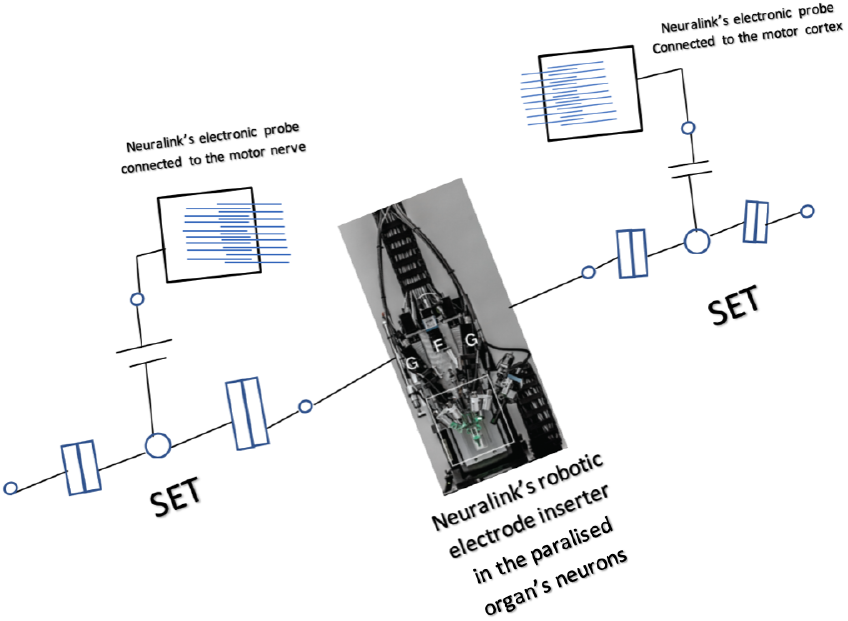
Prototype for Paralysis of Degree 1

#### 2. The mechanism

1. Neuralink’s robotic electrode inserter is implanted to be surrounded by the neurons of the paralysed organ. Where it will pick up the electric field of the generated action and turn them into free electric impulses.
2. The motor data be uploaded on the generated electric impulses.
3. Two single-electron transistors (SET) which are connected; one at each end of the surface of the Neuralink’s robotic electrode inserter, will induce a Cooper pair that allows a pair of electrons to flow freely in superconductors.
4. Two entangled currents will be created as two entangled action potentials, one will be transmitted to the brain by Neuralink’s electronic probe, which is implanted to be connected to the motor cortex, and the second will be transmitted to the motor nerve of the paralysed organ in the same mechanism.
5. The brain will process the data as diagonal matrices of eigenvalues. The data in the motor neuron will be processed as self-adjoint matrices simultaneously. Just then, the paralysed of degree 1 will gain processed motor skill.

### F. A prototype for Paralysis of Degree 2

A paralysed of degree 2 has a deficiency in the entangled Brain-Effector system

#### 1. The components

1. Neuralink’s robotic electrode inserter.
2. Single-electron transistors (SET) which induce a Cooper pair that allows a pair of electrons to flow freely in superconductors. It is clear that we can include the single-electron transistor (SET) at the surface of the Neuralink’s robotic electrode inserter.
3. Neuralink’s electronic probes.
4. Artificial intelligence robot which is able to process the motor data as diagonal matrices of eigenvalues.
5. A bionic organ.

#### 2. The mechanism

1. Neuralink’s robotic electrode inserter is implanted to be surrounded by the neurons of the frontal lobe. Where it will pick up the electric field of the generated action and turn them into free electric impulses. The frontal lobe will play the guidance role instead of the motor neurons and create the motor data as critical thinking data.
2. The critical thinking data be uploaded on the generated electric impulses.
3. Two single-electron transistors (SET) which are connected; one at each end of the surface of the Neuralink’s robotic electrode inserter, will induce a Cooper pair that allows a pair of electrons to flow freely in superconductors.
4. Two entangled currents will be created as two entangled action potentials, one will be transmitted out of the brain to the artificial intelligence robot, and the second will be transmitted to bionic organ, which is connected directly to the paralysed organ.
5. The Artificial intelligence robot will process the data as diagonal matrices of eigenvalues of motor data. Simultaneously, the data will be processed as final solutions; self-adjoint matrices, in the bionic organ. Just then, the paralysed organ of degree 2 will be able to perform the gained motor skill by the connected bionic arm.

**FIG. 12.**
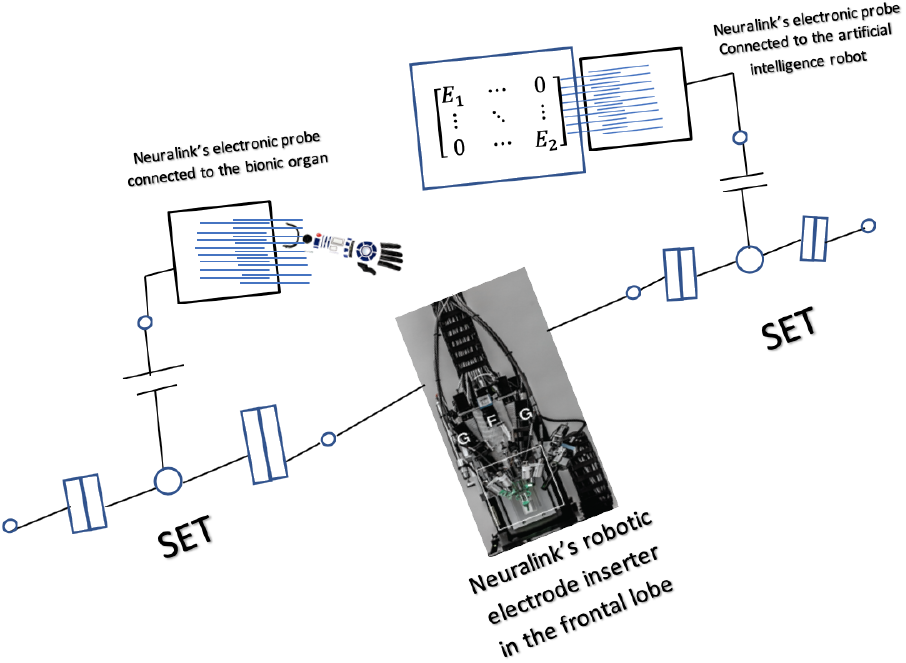
Prototype for Paralysis of Degree 2

### G. A prototype for Alzheimer’s disease

According to the Cognition theory and the eighth result, Alzheimer’s disease can be defined as lack of entanglement between the generated action potentials. Therefore, instead of keeping the sensory action potential as a pattern in the memory lobe, it will do the role of the motor action potential^4,19^.

#### 1. The components

1. Neuralink’s robotic electrode inserter.
2. Single-electron transistors (SET) which induce a Cooper pair that allows a pair of electrons to flow freely in superconductors. It is clear that we can include the single-electron transistor (SET) at the surface of the Neuralink’s robotic electrode inserter.
3. Neuralink’s electronic probes which transmit data to the brain and the effector accurately.

#### 2. The mechanism

1. Neuralink’s robotic electrode inserter is implanted to be surrounded by the frontal lobe neurons. Where it will pick up the electric field of the generated action and turn them into free electric impulses.
2. The pattern data be uploaded on the generated electric impulses.
3. Two single-electron transistors (SET) which are connected; one at each end of the surface of the Neuralink’s robotic electrode inserter, will induce a Cooper pair that allows a pair of electrons to flow freely in superconductors.
4. Two entangled currents will be created as two entangled action potentials, one will be processed at the frontal lobe and the second will be transmitted to the sensory lobe to be kept as a pattern by Neuralink’s electronic probe, which is implanted to be connected to the memory lobe.
5. The frontal lobe will process the data as diagonal matrices of eigenvalues. The data in the memory lobe will be processed as self-adjoint matrices simultaneously. Just then, Alzheimer’s disease patient can remember the kept data again.

**FIG. 13.**
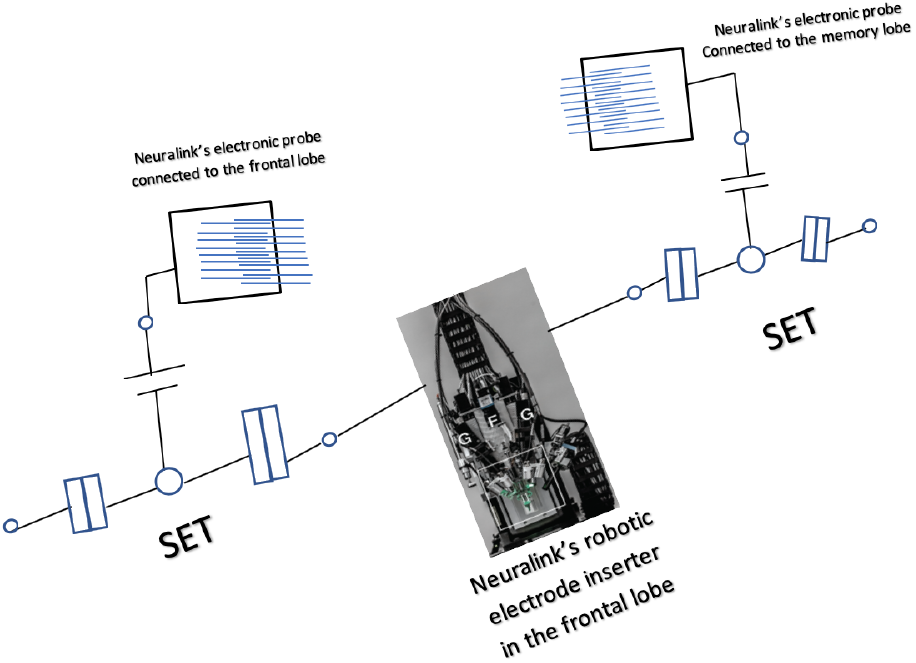
Prototype for Alzheimer’s disease

## CONCLUSION

Human-Brain Artificial-Intelligence Matrix is a new technology that aims to figure out how our neurons and brain work and how to enable them to do their roles properly in case of a deficiency. This technology based on the Cognition theory which can be summarised in two main ideas: First, the action potential arises as two entangled impulses, one in a sensory neuron, and the other in the entangled motor neuron. Next, the state of entanglement between neurons will be replaced with a state of entanglement between the brain and the effector. Second, the brain processes arbitrary data through creation diagonal matrices of discrete eigenvalues, where, the created processed matrix in the brain and the created self-adjoint matrix in the effector are identical. These main results were the scientific basis to propose a general application and classify two main types of deficiency. The first was the entangled Sensory-Motor neurons deficiency which referred to as Degree 1. The second was he entangled Brain-Effector system deficiency which referred as Degree 2. Next, this general idea became more specific to include particular types of deficiency. These types are blindness of degree 1, blindness of degree 2, deafness of degree 1, deafness of degree 2, paralysis of degree 1, paralysis of degree and Alzheimer’s disease. In details, each type of these deficiencies was discussed in three main ideas: the components, the mechanism and the design. Finally, a feasibility study was prepared in three main focuses: cost, employees and duration.

## ACKNOWLEDGMENTS

I would like to thank my family for encouraging me to think critically, work on the scientific field and innovating new ideas and inventions, serve humanity and science. I also would like to thank Dr Moheb for his continual support and for his caregiving during the last three years. Another special thank to Beshoy Mikhael for his tireless efforts on managing the project professionally and presenting all the possible, creative ideas for the sake of the success of the project. Finally, I would like to thank “bioRxiv” for having a very good reputation and thanks to all the great editors for their great efforts. ....

## DATA AVAILABILITY

The data that support the findings of this study are available from the corresponding author upon reasonable request.

## REFERENCES

1 “Sciencedaily, quantum entanglement.”.

2 “Overview of neuron structure and function.”.

3 M. A. L. Tara C. Thiagarajan, “Coherence potentials: Loss-less, all-or-none network events in the cortex.” (2010).

4 D. Robson, “Brain ‘entanglement’ could explain memories.” (2010).

5 C. J. Davisson and L. H. Germer, (1928).

6 “How neurons communicate,” (2018).

7 C. Erkal’s, “Electric potential energy (u) and electric potential (v).”.

8 A. I. M. Rae, “Quantum mechanics, sixth edition,”.

9 “Hyperphysics, the hamiltonian.”.

10 “Porter, f. linear differential equations.”.

11 A. Zettl, “Sturm-liouville theory.”.

12 I. I. O. Physics, “Entanglement: The nature of quantum correlations.” ().

13 I. I. O. Physics, “Entangled spin 1/2 particle pairs and hidden variables.” ().

14 I. I. O. Physics, “Entangled spin 1/2 particle pairs versus an elementary hidden variable theory.” ().

15 S. Csonka, “Cooper pairs as a source of entanglement.” (2010).

16 ORCAM, “What do blind people see?” (2018).

17 E. M. Neuralink, “An integrated brain-machine interface platform with thousands of channels.” (2019).

18 C. Clinic, “Paralysis,”.

19 F. David Walker Dr., Tavares, “The quantum theory of entanglement and alzheimer’s.” (2019).

